# A multimodal fMRI dataset unifying naturalistic processes with a rich array of experimental tasks

**DOI:** 10.1101/2024.06.21.599974

**Authors:** Heejung Jung, Maryam Amini, Bethany J. Hunt, Eilis I. Murphy, Patrick Sadil, Yaroslav O. Halchenko, Bogdan Petre, Zizhuang Miao, Philip A. Kragel, Xiaochun Han, Mickela O. Heilicher, Michael Sun, Owen G. Collins, Martin A. Lindquist, Tor D. Wager

## Abstract

Cognitive neuroscience has advanced significantly due to the availability of openly shared datasets. Large sample sizes, large amounts of data per person, and diversity in tasks and data types are all desirable, but are difficult to achieve in a single dataset. Here, we present an open dataset with N = 101 participants and 6 hours of scanning per participant, with 6 multifaceted cognitive tasks including 2 hours of naturalistic movie viewing. This dataset’s combination of ample sample size, extensive data per participant, more than 600 iso hours worth of data, and a wide range of experimental conditions — including cognitive, affective, social, and somatic/interoceptive tasks — positions it uniquely for probing important questions in cognitive neuroscience.

## Background & Summary

Neuroimaging has significantly advanced our understanding of the dynamics of human mind, specifically at the macroscale level. This progress – which nicely complements human behavioral studies and animal models of micro-meso scale neuroscience – is driven by developments in analytic methods, the high quality of imaging data, the increase in sharing practices of open datasets, and extensive data collection efforts. In large scale studies, two primary approaches are employed: either sampling a large number of participants to enhance statistical power for group average analyses (increase between-subject sampling) or collecting extended hours within participants to better detect specific processes (increase within-subject sampling;^1^). Here, leveraging both between-subject and within-subject sampling, we introduce a dataset that adeptly balances a large sample size of 101 participants with 6 hours worth of neuroimaging data per participant. This dataset is positioned to deepen our understanding of individual brain function, and also highlight individual differences of cognitive processing, given the statistical power that this dataset offers.

Another line of consideration in data collection has been the choice between naturalistic and experimental paradigms. Recently, naturalistic stimuli have proven invaluable in bringing about scientific advances in the neuroimaging community^2–4^, providing rich contexts for exploring a wide range of features from perceptional, situational, to abstract conceptual levels. They offer ecological validity, enabling deeper insights into sensory perception or abstract processing, as the meaning of a stimulus unfolds in a dynamic context in the real world. For example, dynamic stimuli elicit greater activation in the fusiform face area (FFA) compared to static images, uncovering the fact that the FFA may be combining object oriented information with motion^5^. Similarly, dynamic presentations enhance emotion identification^6^, emphasizing the importance of presenting ecologically valid stimuli, as different presentation styles of stimuli may lead to different conclusions. However, it is important to note the inherent challenges of these naturalistic approaches, as they often yield noisier parameters and require complex modeling approaches compared to traditional experimental designs.

For these reasons, experimental designs are indeed crucial in understanding human brain function. In fact, causal relationships of treatment effects are only guaranteed through manipulation and randomization of variables^7–10^. A well-defined parameter space in experiments allows for testing specific hypotheses, developing precise models, and increasing efficiency and statistical power. The rigor and precision of experimental designs are precisely why they are considered the gold standard in biomedical science and why regulatory agencies require randomized controlled trials for drug approvals. These designs effectively isolate the variables of interest and minimize the influence of confound variables. As is well known, fMRI data, specifically the Blood Oxygen Level-Dependent (BOLD) response, is delayed and cumulative in nature as opposed to neural activity, thereby complicating signal attribution to a variable of interest. However, extensive research on the signal profile and optimized experimental designs^11–13^ have mitigated these limitations, enhancing the accuracy of interpretations of the defined parameter space, driving advancements in human neuroimaging for decades.

By integrating both naturalistic and experimental stimuli, we are able to uncover common neural properties that bridge the gap between ecologically valid contexts and experimentally controlled conditions. This integrated approach in past studies has led to significant insights. For example, in an fMRI study on action perception, researchers find several visual processing regions that are consistently activated for both static and dynamic conditions, indicating invariance to form or presentation modality^14^. Similarly, a study on numerical processing using electrocorticography^15^ revealed that the intraparietal sulcus, active in numeracy tasks in experimental settings, are also active during social conversations in naturalistic settings involving numerical concepts. This combined approach not only identifies the homology between the two different settings, but also highlights the subtle distinctions inherent in each process, offering a more holistic understanding of human brain functions.

Harnessing the power of both approaches, our dataset includes 120 minutes of naturalistic movie data and audio narratives, complemented by subjective ratings from participants, and a range of experimental tasks, including somatic, social, cognitive, and affective experimental conditions. Such an approach leverages both ecological validity and statistical power. We envision this dual approach to serve as a versatile tool for probing brain function.

One primary use case of this dataset would be to develop functional alignment techniques to address the challenges of individual differences in neuroimaging. Alignment methods^16^, such as hyperalignment, connectivity alignment, and shared response models, provide solutions to narrow the gap across individuals and help uncover group level neural processes. Since utilizing functional training data from the same individuals are essential for implementing these techniques effectively, the 120 minutes of movie data provides an ideal test environment. Additionally, the inclusion of experimental tasks allows for the rigorous testing and comparison between different functional alignment methods and their validation, enhancing our understanding of both individual and group-level neural processes.

## Methods

### Participants

This dataset includes 101 adult participants (mean ± s.d. age: 24.7 ± 5.5 years; 69 males, 45 females, 2 others). Data were collected from December 2020 to July 2022 at the Dartmouth Brain Imaging Center. Participants were healthy individuals, with normal or corrected-to-normal vision and hearing, no recent psychiatric or neurological diagnoses within the past six months, no MRI contraindications, and no chronic pain. We determined eligibility via a general health questionnaire, a pain safety screening form, and an MRI safety screening form. Additionally, individuals with self-reported chronic pain who anticipated discomfort while lying in an MRI scanner were not enrolled. Participants were recruited from the area of New Hampshire and Vermont as well as the Dartmouth college student body. The institutional review board of Dartmouth College approved the study, and all participants provided written consent. All participants were right handed.

### Overview of experimental procedures

This dataset includes multimodal data from four in-person neuroimaging sessions and an at-home questionnaire survey session. Upon arrival for each in-person neuroimaging session, participants were invited to a separate behavioral testing facility to complete a behavioral informational session, where they signed consent forms and then had scripted task instructions read to them, accompanied by visual aids of upcoming tasks. Participants then completed a behavioral practice task on a computer to reinforce their understanding of the task. After the behavioral information session, participants were invited to the MRI scanning facility, in which they completed experimental tasks with concurrent fMRI and physiological recordings of skin conductance and photoplethysmography (Fig. 1; see Table 1 for imaging acquisition parameters). Participants completed at-home surveys prior to the first incoming scan session. These surveys primarily assessed general psychosocial tendencies, with the aim to link them with behavioral, physiological, and neural responses observed during the neuroimaging experiments (Table 2).

**Figure 1.**
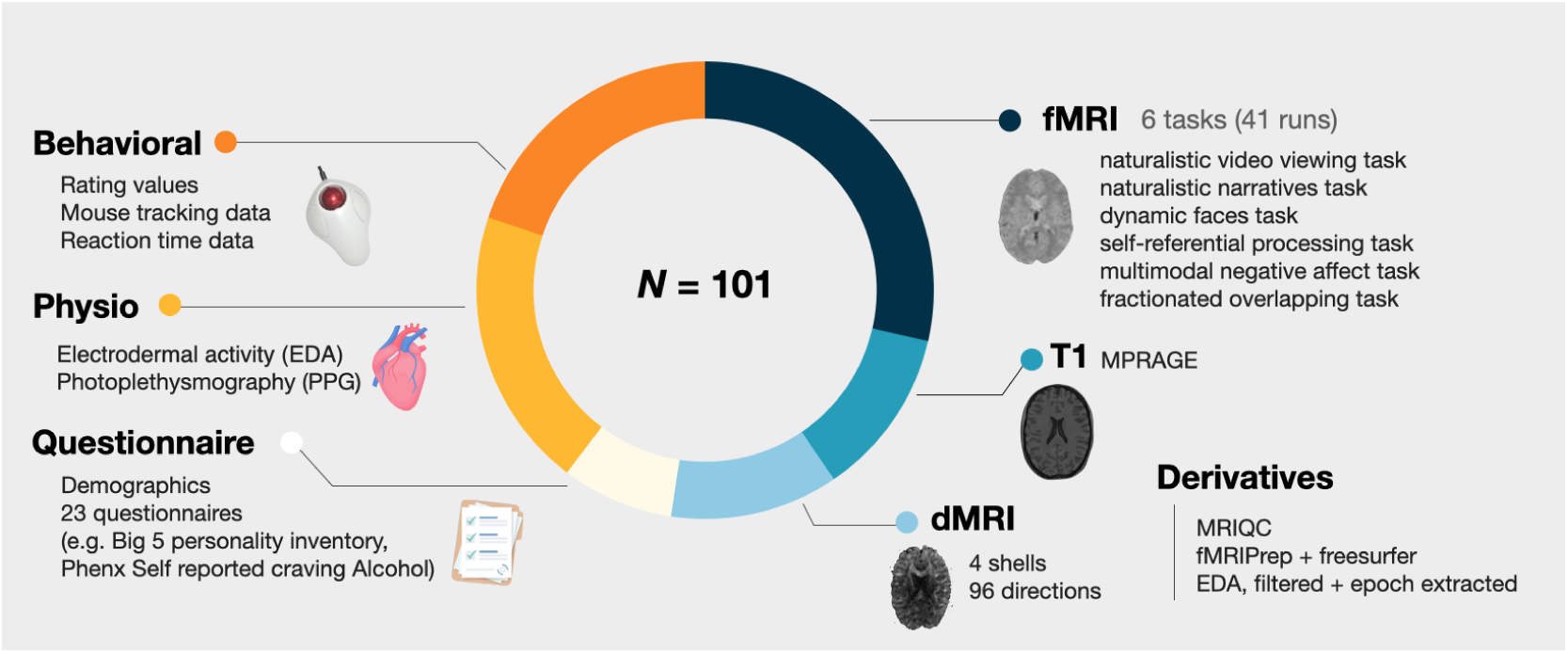
Overview of acquired data. Dataset includes N = 101 participants worth of data, including (*clockwise*): 1) functional BOLD echo-planar imaging with cognitive tasks (TR = 460 msec, MB = 8), 2) a T1-weighted anatomical scan, 3) a multi-shell diffusion weighted MRI (dMRI) scan, 4) a battery of questionnaires prior to scanning, 5) physiological data collected during scanning, and 4) behavioral data collected during scanning.

**Table 1.** Description of imaging acquisition parameters.

| Features | Main Task | T1w | DWI | Distortion Correction |
| --- | --- | --- | --- | --- |
| Resolution [mm] | 2.7 (iso) | 0.8 (iso) | 1.7 (iso) | 2.7 (iso) |
| # Volumes | Varies per task | 1 | 1 | 2 |
| FOV [mm] | 220 | 256 | 240 | 220 |
| Matrix Size | 82×82 | 320×320 | 130×130 | 82×82 |
| TR [s] | 0.46 | 2 | 4.11 | 7.22 |
| TE [ms] | 27.20 | 2.11 | 88.40 | 73.00 |
| Flip Angle [degree] | 44° | 8° | 90° | 90° |
| Slice Orientation | Transversal | Sagittal | Transversal | Transversal |
| Phase Encoding Direction | A >> P | A >> P | A >> P | A >> P |
| Number of Slices | 56 | 224 | 81 | 56 |
| Slice Thickness [mm] | 2.7 | 0.8 | 1.7 | 2.7 |
| Distance Factor [%] | 0 | 50 | 0 | 0 |
| Order of Slice Acquisition | Interleaved | Interleaved | Interleaved | Interleaved |
| iPAT mode | none | GRAPPA | none | none |
| Multiband Acceleration Factor | 8 | 3 | 3 | 1 |
| Bandwidth [Hz/px] | 3048 | 240 | 1700 | 3048 |

**Table 2.**
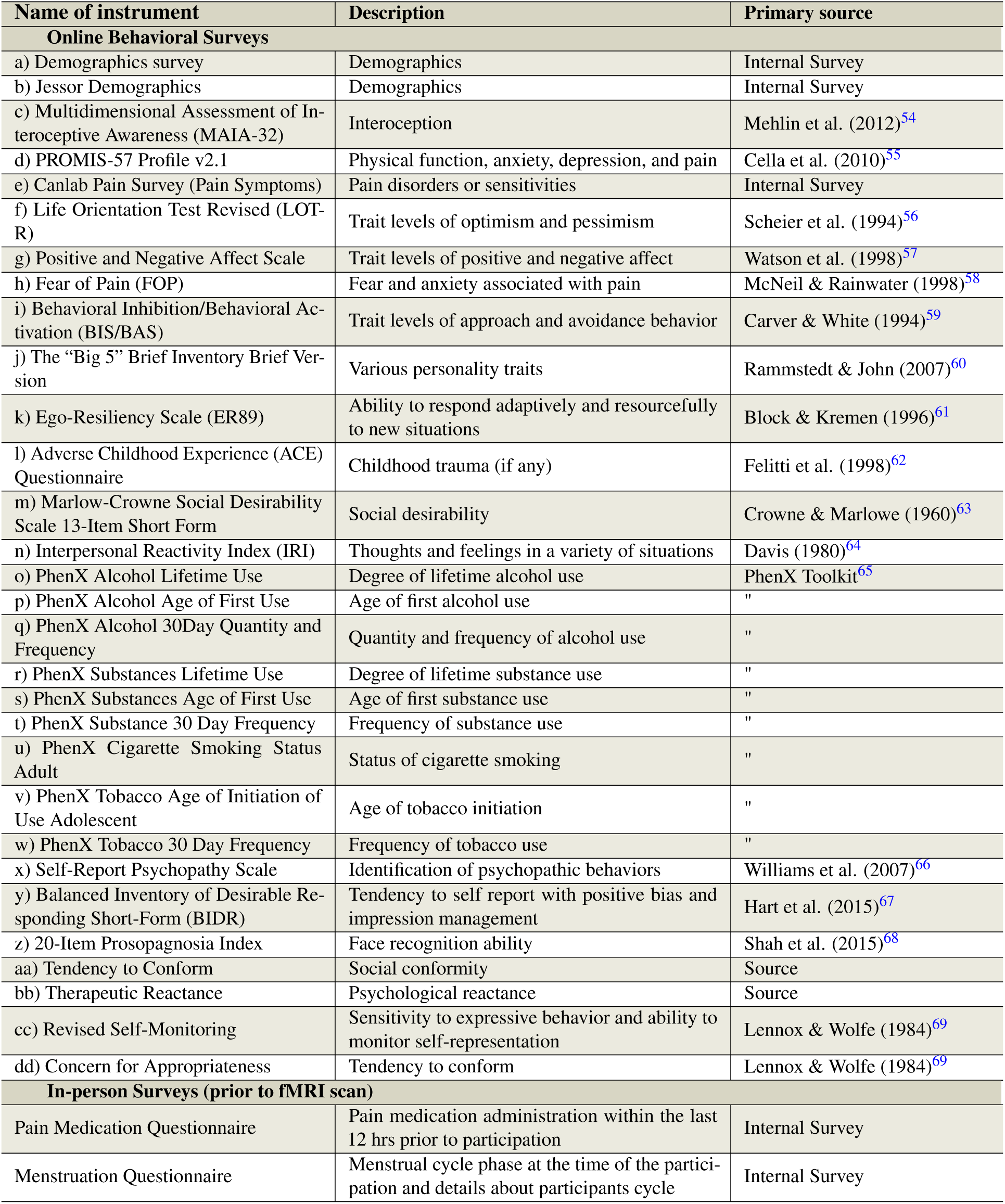
Description of questionnaires. Participants completed at-home surveys prior to initial neuroimaging sessions. “Name of instrument” column indicates the short hand title of each questionnaire. “Description” indicates a short summary of the aim of each questionnaire and “Primary source” indicates the reference for each questionnaire.

### Overview of neuroimaging tasks

Each neuroimaging session encompassed a variety of tasks, each designed to probe different cognitive domains (illustrated in Fig. 2: “task description”). For example, in `task-alignvideo`, participants watch naturalistic videos and rate their emotional responses. Therefore, this task includes multimodal stimuli, incorporating both visual and auditory elements, and incorporates affective and social domains. Conversely, `task-faces` specifically focuses on face processing, involving visual recognition and interpretation of affective facial expressions. Detailed descriptions of each task are provided in subsequent subsections, denoted as “`task-`”. Visual representations of individual trials within each task are depicted in Fig. 3. While analysis of interest may vary across researchers, we provide a detailed overview of the key contrasts inherent to each task (Table 3). This includes a description of the primary cognitive processes, or “canonical” contrasts, which serve for understanding the core aspects of each tasks’ design and its analytical focus.

**Figure 2.**
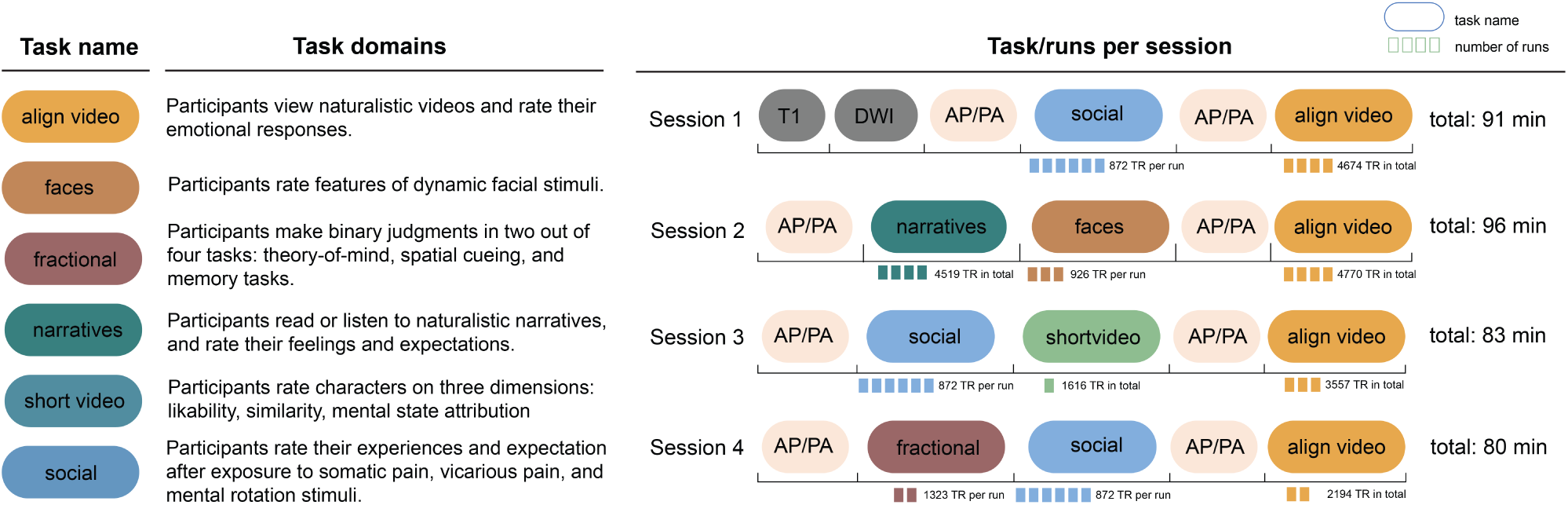
Data acquisition layout. Each neuroimaging session included task data from multiple cognitive domains, as demonstrated in the left panel. As participants completed multiple tasks across four sessions, sequence of each task and number of runs per task are illustrated in the right panel. Session 1 consisted of one anatomical T1-weighted scan, multi-shell diffusion weighted image, AP/PA image for distortion correction, followed by six EPI runs of the `social` task, another AP/PA image for distortion correction, followed by four EPI runs of `alignvideo` task. Session 2 started with a AP/PA image, followed by four narrative EPI runs, three faces EPI runs, followed by AP/PA run, and four EPI runs of the video task. Session 3 consisted of an AP/PA run, followed by six EPI runs of the social task, one EPI run of the `shortvideo` task, AP/PA run, and ended with three EPI runs of the align video task. Session 4 started with an AP/PA run, followed by two EPI runs of the fractional task, six EPI runs of the social task, another AP/PA run, followed by two EPI runs of the align video task.

**Figure 3.**
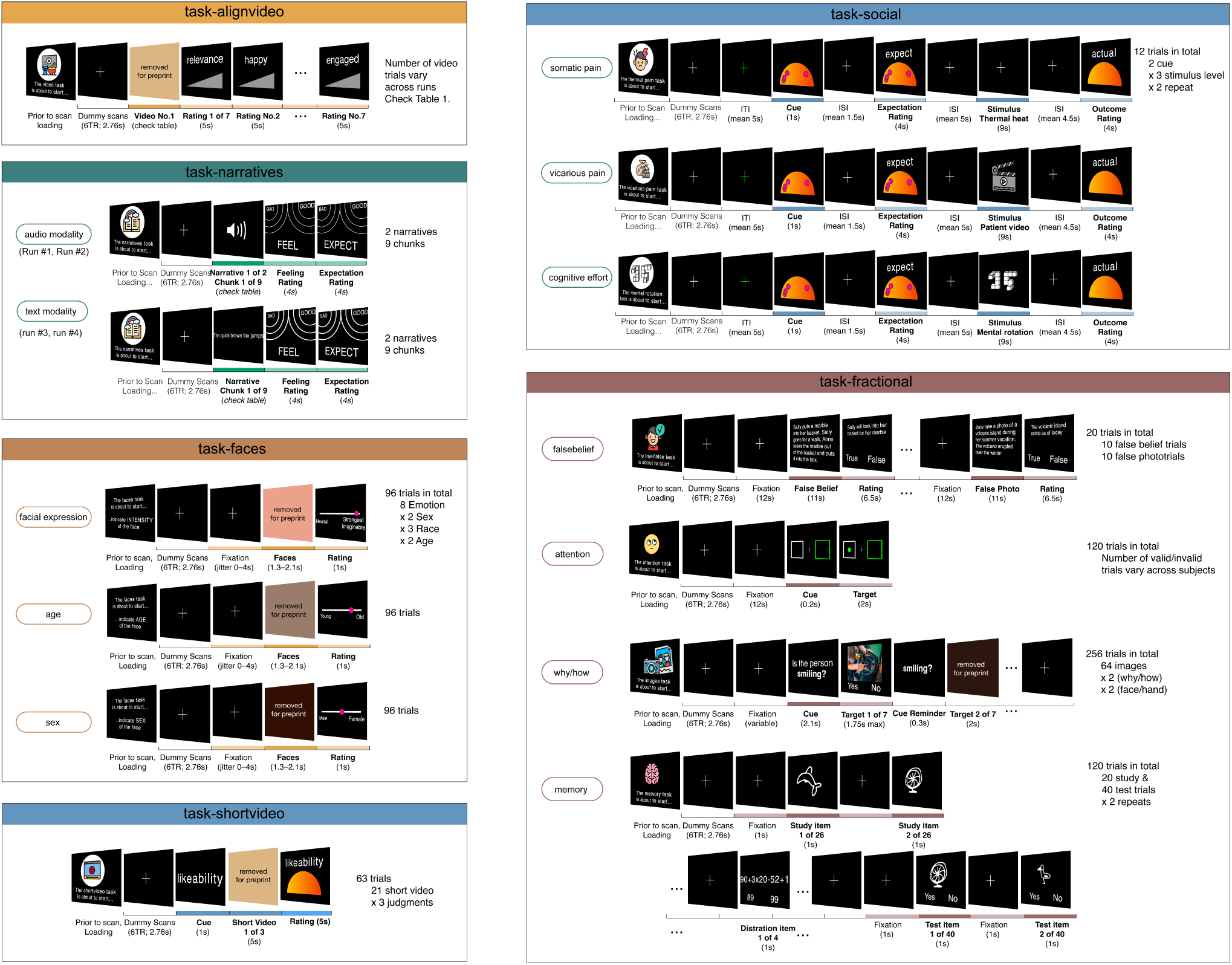
Depiction of task structure and its variations across runs. Each of the six panels refers to a task. Within each panel, “trials” are described, which are fundamental units of a run, designed to capture a specific cognitive process. Trials are repeated within runs. The number of these repetitions, which may vary across tasks, is indicated on the right side of the panel. For tasks employing a factorial design, — specifically, `task-faces, task-shortvideo, task-social, task-falsebelief, task-attention, task-whyhow, task-memory` — the experimental factors are also listed on the right. On the left side of each panel, we note any changes in rating or stimulus modality across runs. For example, `task-narratives` encompassed two distinct stimulus modalities: audio and text narratives, delivered in separate runs. In a similar vein, the `task-social` consisted of several distinct runs, each dedicated to a different domain: somatic pain, vicarious pain, and cognitive effort. Stimuli corresponding to each domain were presented exclusively within their respective runs. Lastly, `task-faces` included three runs, each requiring participants to judge faces based on different dimensions, such as sex and age respectively. These variations in modality and rating type across runs are indicated in bubbles on the left side of each panel.

**Table 3.**
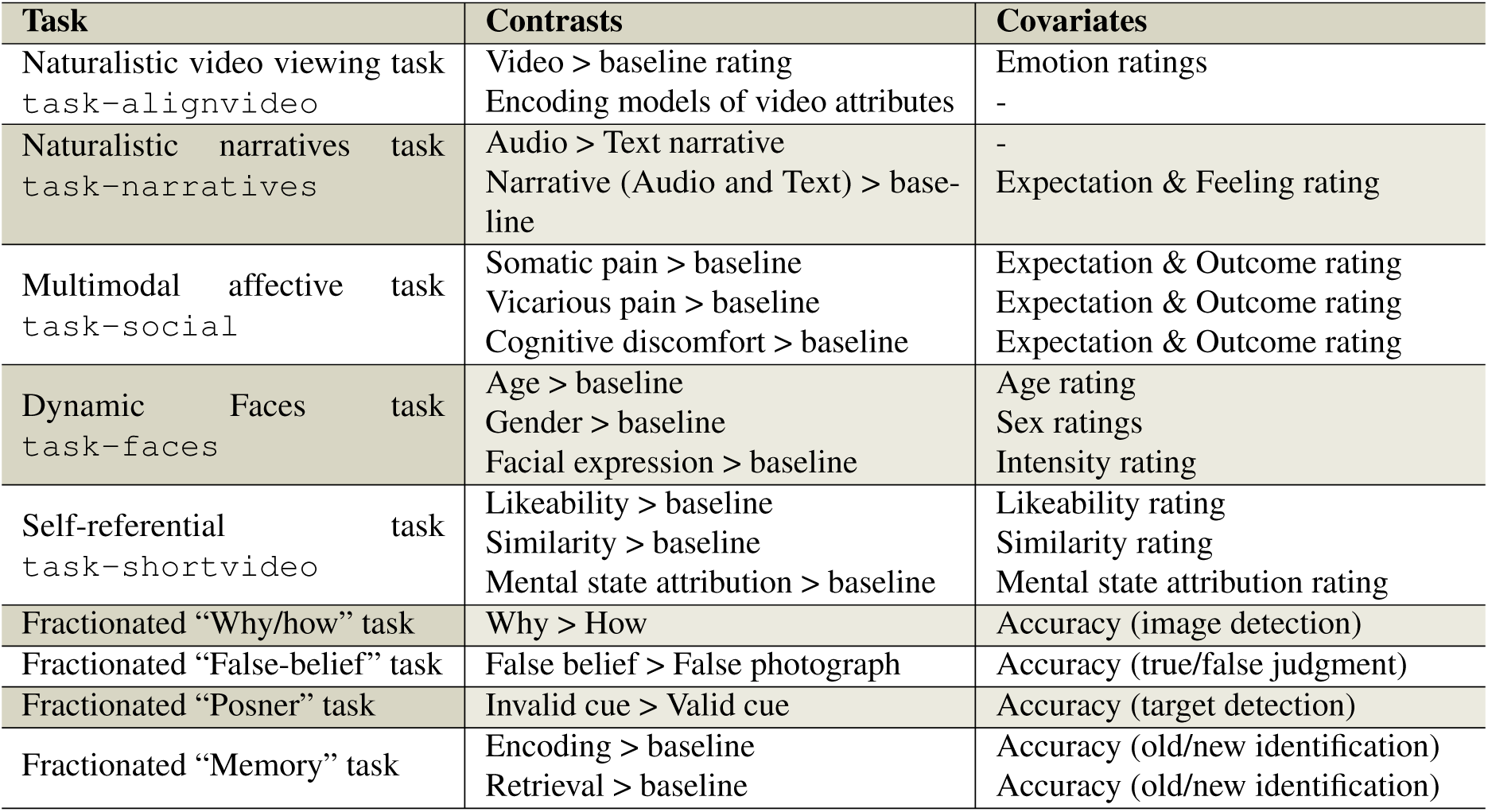
Description of canonical neuroimaging contrasts per task. To facilitate exploration of the dataset, we highlight the primary processes associated with each task by outlining the canonical contrasts each task (listed in “Task” and “Contrasts” column). Additionally, “Covariates” indicate the participant ratings collected during each trial. These covariates could be integrated into within-subject models or included as between-subject individual difference measures after further computation. For example, the Narratives task includes contrasts such as 1) audio vs. text modality comparison, 2) narrative vs. baseline comparison, and 3) the incorporation of behavioral ratings as covariates.

| Task | Contrasts | Covariates |
| --- | --- | --- |
| Naturalistic video viewing task<br>task-alignvideo | Video > baseline rating<br>Encoding models of video attributes | Emotion ratings<br>- |
| Naturalistic narratives task<br>task-narratives | Audio > Text narrative<br>Narrative (Audio and Text) > baseline | -<br>Expectation & Feeling rating |
| Multimodal affective task<br>task-social | Somatic pain > baseline<br>Vicarious pain > baseline<br>Cognitive discomfort > baseline | Expectation & Outcome rating<br>Expectation & Outcome rating<br>Expectation & Outcome rating |
| Dynamic Faces task<br>task-faces | Age > baseline<br>Gender > baseline<br>Facial expression > baseline | Age rating<br>Sex ratings<br>Intensity rating |
| Self-referential task<br>task-shortvideo | Likeability > baseline<br>Similarity > baseline<br>Mental state attribution > baseline | Likeability rating<br>Similarity rating<br>Mental state attribution rating |
| Fractionated “Why/how” task | Why > How | Accuracy (image detection) |
| Fractionated “False-belief” task | False belief > False photograph | Accuracy (true/false judgment) |
| Fractionated “Posner” task | Invalid cue > Valid cue | Accuracy (target detection) |
| Fractionated “Memory” task | Encoding > baseline<br>Retrieval > baseline | Accuracy (old/new identification)<br>Accuracy (old/new identification) |

### Specific tasks, stimuli and rating description

#### Generalized Labeled Magnitude Scale (gLMS)

A challenge in scientific studies of human behavior is the subjectivity of outcome measures, making ratings often non-comparable across studies. For this dataset, we used a modified semi-circular-shaped generalized labeled magnitude scale (gLMS^17^), which allows for magnitude matching across participants, originating from gustatory research and psychophysics. Several tasks in the current dataset, including the Multimodal negative affect task and Self-referential task, use elements of the gLMS. The scale is intended to measure subjective sensory experiences quantifiably and comparably across different sensory/cognitive domains, allowing for comparisons both between and within participants. Labels on the scale range from “No sensation” (0°), “Barely detectable” (3°), “Weak” (10°), “Moderate” (29°), “Strong” (64°), “Very Strong” (98°) to “Strongest sensation of any kind” (180°). To mitigate scale usage bias, the scale was adapted from a vertical linear scale to a semicircular presentation, in which reported ratings are designed to be equidistant from the cursor starting point. The scale and associated tasks were presented on a desktop computer during the pre-scan behavioral instruction sessions and in-scanner fMRI tasks. The scale was adapted to each task by modifying the label of the maximum magnitude marker within the context of the task’s measures.

#### Naturalistic video viewing task “task-alignvideo”

##### Task procedures

A naturalistic video-viewing task was administered across all four sessions. Each session consisted of multiple runs (see Fig. 2 for run composition per session); each run consisted of multiple video trials. Each trial consisted of two epochs of watching a video and rating seven emotion categories. First, participants watched a series of emotionally salient videos (“video”) and next, were intermittently prompted to provide ratings (“rating”) about how the video made them feel in relation to seven affective domains: 1) personal relevance, 2) happy, 3) sad, 4) afraid, 5) disgusted, 6) warm and tender, and 7) engaged. Participants were given 5 seconds to make each of the seven ratings; the sequence of these ratings were kept constant. A variety of 49 unique videos were presented to participants, ranging from 20 seconds to 5 min 39 seconds in duration, amounting to 86 minutes and 9 seconds across 4 sessions (See Table 4 for detailed breakdown of video duration, order and, number of TRs). Each video was played once, with no repetitions. The sequence of the videos were identical across participants, purposefully designed for functional alignment purposes.

**Table 4.**
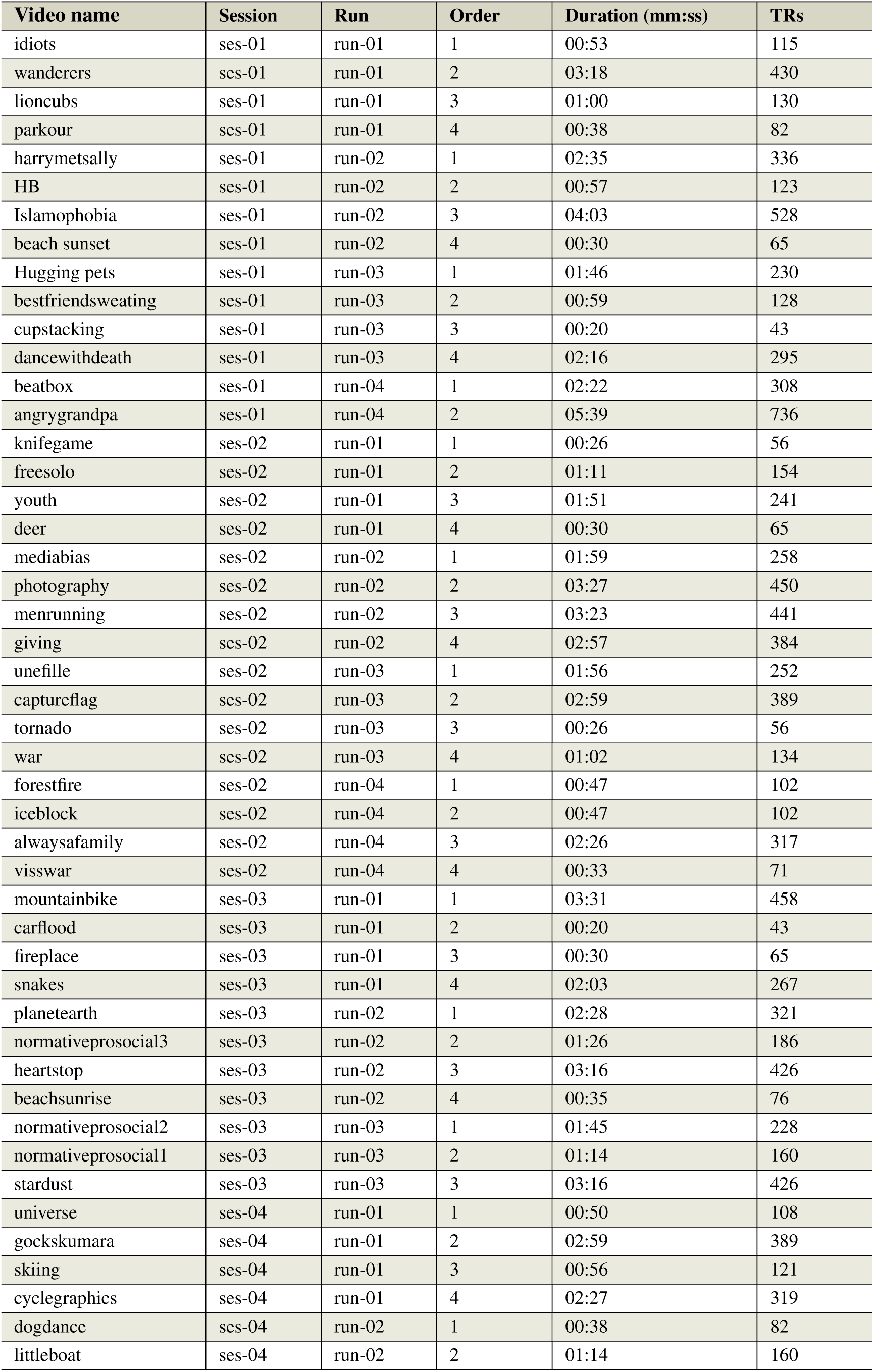
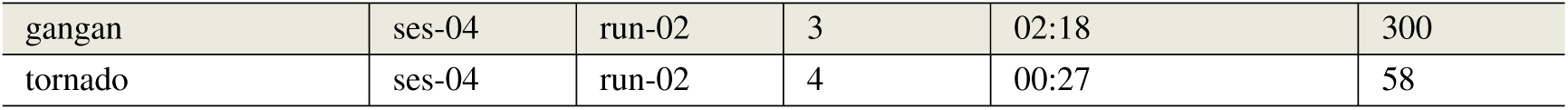
Overview of the naturalistic videos used in the Align video task. Each row represents a video that was presented during one trial. The column “Video” indicates the video filename. “Session” and “Run” indicates the session and run in which the videos were presented “Order” indicates the order of the video played in each run. “Duration (mm:ss)” indicates the length of the video. “TRs” indicates the number of TRs, i.e. 0.46 seconds, that the video spans. Each video file follows the naming convention: “{Session}_{Run}_order-{Order}_content-{Videonames}.mp4”, shared in the “stimuli/task-alignvideo” folder.

| Video name | Session | Run | Order | Duration (mm:ss) | TRs |
| --- | --- | --- | --- | --- | --- |
| idiots | ses-01 | run-01 | 1 | 00:53 | 115 |
| wanderers | ses-01 | run-01 | 2 | 03:18 | 430 |
| lioncubs | ses-01 | run-01 | 3 | 01:00 | 130 |
| parkour | ses-01 | run-01 | 4 | 00:38 | 82 |
| harrymetsally | ses-01 | run-02 | 1 | 02:35 | 336 |
| HB | ses-01 | run-02 | 2 | 00:57 | 123 |
| Islamophobia | ses-01 | run-02 | 3 | 04:03 | 528 |
| beach sunset | ses-01 | run-02 | 4 | 00:30 | 65 |
| Hugging pets | ses-01 | run-03 | 1 | 01:46 | 230 |
| bestfriendsweating | ses-01 | run-03 | 2 | 00:59 | 128 |
| cupstacking | ses-01 | run-03 | 3 | 00:20 | 43 |
| dancewithdeath | ses-01 | run-03 | 4 | 02:16 | 295 |
| beatbox | ses-01 | run-04 | 1 | 02:22 | 308 |
| angrygrandpa | ses-01 | run-04 | 2 | 05:39 | 736 |
| knifegame | ses-02 | run-01 | 1 | 00:26 | 56 |
| freesolo | ses-02 | run-01 | 2 | 01:11 | 154 |
| youth | ses-02 | run-01 | 3 | 01:51 | 241 |
| deer | ses-02 | run-01 | 4 | 00:30 | 65 |
| mediabias | ses-02 | run-02 | 1 | 01:59 | 258 |
| photography | ses-02 | run-02 | 2 | 03:27 | 450 |
| menrunning | ses-02 | run-02 | 3 | 03:23 | 441 |
| giving | ses-02 | run-02 | 4 | 02:57 | 384 |
| unefille | ses-02 | run-03 | 1 | 01:56 | 252 |
| captureflag | ses-02 | run-03 | 2 | 02:59 | 389 |
| tornado | ses-02 | run-03 | 3 | 00:26 | 56 |
| war | ses-02 | run-03 | 4 | 01:02 | 134 |
| forestfire | ses-02 | run-04 | 1 | 00:47 | 102 |
| iceblock | ses-02 | run-04 | 2 | 00:47 | 102 |
| alwaysafamily | ses-02 | run-04 | 3 | 02:26 | 317 |
| visswar | ses-02 | run-04 | 4 | 00:33 | 71 |
| mountainbike | ses-03 | run-01 | 1 | 03:31 | 458 |
| carflood | ses-03 | run-01 | 2 | 00:20 | 43 |
| fireplace | ses-03 | run-01 | 3 | 00:30 | 65 |
| snakes | ses-03 | run-01 | 4 | 02:03 | 267 |
| planetearth | ses-03 | run-02 | 1 | 02:28 | 321 |
| normativeprosocial3 | ses-03 | run-02 | 2 | 01:26 | 186 |
| heartstop | ses-03 | run-02 | 3 | 03:16 | 426 |
| beachsunrise | ses-03 | run-02 | 4 | 00:35 | 76 |
| normativeprosocial2 | ses-03 | run-03 | 1 | 01:45 | 228 |
| normativeprosocial1 | ses-03 | run-03 | 2 | 01:14 | 160 |
| stardust | ses-03 | run-03 | 3 | 03:16 | 426 |
| universe | ses-04 | run-01 | 1 | 00:50 | 108 |
| gockskumara | ses-04 | run-01 | 2 | 02:59 | 389 |
| skiing | ses-04 | run-01 | 3 | 00:56 | 121 |
| cyclegraphics | ses-04 | run-01 | 4 | 02:27 | 319 |
| dogdance | ses-04 | run-02 | 1 | 00:38 | 82 |
| littleboat | ses-04 | run-02 | 2 | 01:14 | 160 |
| gangan | ses-04 | run-02 | 3 | 02:18 | 300 |
| tornado | ses-04 | run-02 | 4 | 00:27 | 58 |

##### Stimuli and scales

To submit affective ratings of videos, the participants were presented with a continuous, linear scale ranging from “Barely at all” to “Strongest imaginable”. Practice ratings were provided using a computer mouse, and experimental ratings were provided using an MR-compatible trackball.

#### Multimodal negative affect task “task-social”

##### Task procedures

Participants performed three different task domains: somatic pain (“pain”), vicarious pain (“vicarious”), and mental rotation (“cognitive”). Each task was designed as a 2 cue (high/low) x 3 stimulus intensity (high/med/low) factorial design. The three tasks were conducted repeatedly on average, one week apart across three sessions (`ses-01, ses-03, ses-04`). Each trial consisted of four events: first, participants passively viewed a presentation of a high or low social cue, consisting of data points that participants believed indicated other people’s ratings for that stimulus presented for 1 second on screen (“cue”); second, participants provided ratings of their expectations on the upcoming stimulus intensity on a gLMS scale for a total duration of 4 seconds overlaid with the cue image (“expectancy rating”); third, participants passively received/viewed experimentally delivered stimuli for each of the mental rotation, vicarious pain, and somatic pain tasks for 5 seconds each (“stimulus”); lastly, participants provided ratings on their subjective experience of cognitive effort, vicarious pain, or somatic pain for 4 seconds (“outcome rating”). In total, each run was designed to last 6 minutes and 41 seconds, i.e., 872 TRs. The task was administered in `ses-01`, `ses-03`, and `ses-04`.

##### Stimuli and scales

Stimuli for the somatic pain task were thermal heat stimuli, administered using a TSA2 system (Medoc) with a 16-mm Peltier contact thermode, delivered to the glaborous skin of the ventral surface of the left forearm. Stimuli at three stimulus intensity levels were delivered (low: 48°C; medium: 49°C; high: 50°C), for a total duration of 9 seconds with a 5 second plateau. Baseline temperature was 32°C. Stimuli for the vicarious task were videos of patients in pain, selected from the UNBC-McMaster shoulder pain expression archive database^18^ and categorized into three stimulus intensity levels using the pre-normed ratings (self-reported pain rating and observer estimated-pain rating) provided from the dataset. Stimuli for the mental rotation task were images from the Ganis & Kievet^19^ dataset, selecting images that were rotated 50, 100, 150 ° to account for three intensity levels.

#### Naturalistic narratives task “task-narratives”

##### Task procedures

Participants were instructed to read or listen to 8 different narratives while in the scanner, with each narrative chunked into 9 clips. Each trial started with a fixation period with jittered durations between 2 and 8 seconds (“fixation”). During the narrative epoch, audio or text clips were presented (“audio” or “text”). Participants were then prompted to rate how they felt about the narrative (“feeling”) and what their expectations were for the upcoming narrative on how good or bad the future storyline would be (“expectation”). The entire task consisted of four runs, with two narratives, i.e. 18 narrative clips in each run. In total, each run was designed to last 7 minutes 24 seconds to 9 minutes 57 seconds, i.e., 967, 1098, 1298, 1156 TRs per run. The task was administered in `ses-02`.

##### Stimuli and scales

The narrative contents were manipulated across three distinct timescales: individual situations, the context of those situations, and the overarching full narrative. Each narrative includes a single main character who moves between three different contexts and, in each context, encounters three different situations (characterized by interpersonal relationships and actions). We use Polti’s 36 situations to construct these different situations^20^. The duration of the eight narratives range from 1 minute 36 seconds to 3 minutes 29 seconds (Table 5). The rating scale had two poles of the scale, each labeled “good” and “bad”, presented at the two upper corners of the screen. Ratings can be operationalized as the distance between participants’ clicks and the centers of the two poles. The advantage is that the scale allows for encoding two dimensions of a participants subjective experience, i.e. intensity and valence, instead of presenting a linear scale.

**Table 5.** Overview of the naturalistic narratives used in the Narratives task. The narrative task was operated within a single session, `ses-03`. There were four runs in total, with 8 different narratives. Each row represents a narrative, presented within each run. The “Run” column indicates the run order in which the narrative was presented. “Story” represents the narrative index. “Modality” column indicates the form in which each narrative was presented. “Duration (mm:ss)” indicates the length of the narrative. “TRs” indicates the number of TRs, i.e. 0.46 seconds, that the audio or text-based narrative spans. Each narrative was further divided into nine chunks labeled by situations; each chunk was presented per trial.

| Run # | Story # | Modality | Duration (mm:ss) | TRs | Words |
| --- | --- | --- | --- | --- | --- |
| run-01 | 7 | audio | 02:01 | 263 | 429 |
| run-01 | 8 | audio | 01:36 | 209 | 314 |
| run-02 | 5 | audio | 02:30 | 326 | 511 |
| run-02 | 6 | audio | 02:04 | 269 | 447 |
| run-03 | 3 | text | 03:11 | 415 | 573 |
| run-03 | 4 | text | 03:29 | 454 | 626 |
| run-04 | 1 | text | 02:43 | 355 | 490 |
| run-04 | 2 | text | 02:40 | 348 | 480 |

#### Dynamic faces task “task-faces”

##### Task procedures

Participants were presented with 288 dynamic faces of varying race, age, sex, and facial expressions. The faces task consisted of three runs; in each run, participants were prompted to rate specific dimensions of the face: age, sex, or intensity of the facial expression. The order of the runs were pseudo-randomized based on odd and even numbered participant IDs. Participants were made aware of the rating dimension before the start of each run. In each trial, a brief video featuring a single face, displaying an expression, was played (“faces”), and participants were given 1.875 seconds to rate the face on the corresponding rating: the intensity of the facial expressions, the sex of the face, or the age of the face (“rating”). Responses were either recorded during manual entry or by recording the position of the cursor at the end of the response period. After each rating, a fixation cross was displayed with a jittered duration of 0-4 seconds. Each of the three runs lasted 7 minutes and 7 seconds, (i.e., 914 TRs) and featured 96 facial stimuli. The task was administered within a single session, during `ses-02`. ***Stimuli and scales.*** The stimuli were animated clips of human faces, ranging from 1.3 to 2.1 seconds, varying across four underlying factors: age, sex, race, and facial expression. The faces were either young or old (“age”), male or female (“sex”), Eastern Asian, Western Caucasian, or African (“race”) and exhibited one of eight emotions: happiness, surprise, fear, disgust, anger, sadness, pain, and pleasure (“expression”). The faces were crossed with the four factors. The rating scale was a linear scale with keywords at each end, serving as axes for the rating scale (i.e. young-old, male-female, neutral-strongest imaginable for each age, sex, expression block of trials).

#### Self-referential task “task-shortvideo”

##### Task procedures

The self-referential task is a theory-of-mind task in which participants are prompted to make three different types of assessments about a featured character from a video clip: 1) similarity, 2) likeability, 3) mental state attribution. Participants were familiar with the featured character because the short video clips were pulled from full-length videos that participants had watched in previous sessions, specifically during `task-alignvideo`. Participants rated three types of ratings per video: 1) perceived similarity towards the character (“similarity”), worded as ‘How similar are you to this character?’; 2) likeability towards the character (“likeability”) worded as ‘How much do you like this character?’; and 3) inferring what the character is thinking based on a question prompt, such as ‘Did the character feel in danger?’ or ‘Was the character remembering something?’ (“mental state attribution”). At the beginning of each block, participants were visually presented with one of these three questions prior to watching a set of three character-videos. Subsequently, participants were given a rating cue (e.g. ‘how similar?’) and were shown a five-second video clip of the character. Once the clip ended, participants had five-seconds to respond to the question using a modified version of the semi-circular gLMS rating scale. To ensure participants were clear about which character the experiment referred to, several measures were implemented: 1) The characters were introduced in videos during earlier sessions, 2) participants rewatched these videos during a pre-scan instructional session, and 3) a slideshow was used to highlight the face of the character being discussed, after participants completed rewatching the videos. We asked participants recognition of the characters; videos were replayed if the participants were not able to recognize the said character. In total, each run was designed to last 12 minutes and 23 seconds, i.e., 1616 TRs. The task was administered within a single session, during `ses-03`.

##### Stimuli and scales

Participants were presented with a series of short video clips, ranging from 4.1 to 6.9 seconds (“video”). As mentioned, these video clips were pulled from previous sessions, and edited to single out a character of interest. Immediately following each video clip, participants were prompted to provide character assessments using a modified version of the gLMS (“rating”). The modified gLMS rating scale included the following anchors: Not at all, barely detectable, weak, moderate, high, very high, strongest possible similarity/likability. For mental ease on rating self-referential thoughts on the videos, we cued participants with the type of rating – likeability, similarity, mental state attribution – at the beginning of a video. Afterwards, three trials consecutively asked the same questions. In other words, a rating question was first presented, followed by one video, then followed by a shorter question cue, and this was iteratively done for three consecutive trials.

#### Fractionated overlapping task “task-fractional”

The fractionated task consisted of four subtasks: attention reorienting^21^, memory encoding/retrieval, text-based theory of mind^22^, and image-based theory of mind^23^. The purpose of this task was to select and compare across subtasks that are known to engage the angular gyrus. Each participant was pseudo-randomly assigned to undergo two subtasks, which were counterbalanced across participants. In other words, one set of participants completed the attention reorienting task and image-based theory of mind task, while a different participant completed a text-based theory of mind task and memory/encoding task. This task was completed within one session, during `ses-04`.

#### Fractionated task A: Attention reorienting task “runtype-posner”

##### Task procedures

This subtask was a canonical spatial cueing paradigm^21^, which consists of three phases: fixation phase, cue phase, and target search phase. The goal is to identify and indicate the location of the *target* as quickly as possible. During the fixation phase, participants were presented with a fixation cross and two empty square boxes outlined in white, positioned in the left and right visual fields. This was displayed for a jittered duration, averaging around 2.5 seconds (“fixation”). During the cue phase, one of the boxes was highlighted with a green color for 0.2 seconds (“cue”). During the target search phase, a green circle, the target, appeared in the center of either the left or right square box for 2 seconds (“target”). If the participant responded within this 2-second window, a feedback screen highlighted the box they selected, for 0.5 seconds. Afterwards, the next trial begins, with the initiation of the fixation cross and empty white boxes on the screen. In total, each run was designed to last 10 minutes and 8 seconds, i.e., 1322 TRs.

##### Stimuli and scales

Stimuli were simple geometric shapes, i.e. boxes and circles. No rating scale was used in this subtask. Participants responded to the target using the two buttons of an MR-compatible trackball.

#### Fractionated task B: Memory encoding-retrieval task “runtype-memory”

##### Task procedures

This subtask was designed to be identical to encoding retrieval tasks, which consist of three phases, repeated twice: a memory encoding phase, a memory distraction phase, and a memory recall phase. The memory encoding phase entailed the presentation of a small illustration clip-art presented for 1 second each (“encoding”), with a total of 26 × 2, i.e., 52 images. The memory distraction phase, intended to prohibit active memory maintenance, included a math calculation task with two math problems × 2 (“distraction”), each displayed for 25 seconds followed by a 5 second response epoch. During the memory recall phase, participants were presented with clip-arts one by one, with a total of 40 × 2 images, and were asked whether they had seen each clip-art before or not. Participants then responded “old” or “new” by pressing the appropriate button under each clip-art. Each response phase lasted 2 seconds (“retrieval”). In total, each run was designed to last 10 minutes and 8 seconds, i.e., 1322 TRs. Due to the recency and primary memory effects, the first and last three images in the encoding test are labeled as null trials.

##### Stimuli and scales

Stimuli were image-based drawings of everyday objects or animate beings. No rating scale was used. Participants responded to the target using the two buttons of an MR-compatible trackball, indicating “old” or “new” to the image presented on screen.

#### Fractionated task C: Theory of mind image-based task “runtype-tomspunt”

##### Task procedures

This subtask was derived from Spunt & Adolphs’ theory of mind why/how localizer^23^ to investigate mental state attribution. The experiment focuses on two cognitive aspects of observable actions: mentalizing the implicit mental states driving the actions (“why”) and describing the explicit physical attributes of these actions (“how”). In each trial, participants first saw a prompt (“question”) for 2.1 seconds, followed by a photo (“photo”) with a duration of 1.75 seconds, and then answered yes/no to the prompted question. In total, each run was designed to have 256 images, with 64 images crossed with 2 questions (why/how) and 2 mediums (face/hand), and lasted 10 minutes and 8 seconds, i.e., 1322 TRs.

##### Stimuli and scales

An example of a stimulus includes a photo that depicts an individual looking sideways. One question would ask “Is this person looking away? (how)” and another would ask “Is this person expressing doubt? (why)” The same image was yoked with the 2 (why/how) × 2 (face/hand) combination of questions, which engaged different processes related to physical vs. mental state attribution.

#### Fractionated task D: Theory of mind text-based task “runtype-tomsaxe”

##### Task procedures

This subtask was derived from Dodell-Feder, Dufour, and Saxe’s theory of mind false belief localizer task^22^ to investigate theory of mind processes, i.e., the process of representing and attributing mental states of another agent. Participants read anecdotes of false beliefs and false photographs and answered a true/false questions regarding the stories. In each trial, a fixation cross was presented for 12 seconds (“fixation”), after which participants read an anecdote for 14 seconds (“story”) and responded to a question regarding the anecdote, presented on screen for 10 seconds (“question”). In total, each run was designed to last 10 minutes and 8 seconds, i.e., 1322 TRs.

##### Stimuli and scales

“False belief” stories were narratives where the reader had to represent outdated beliefs of an agent’s latent thought to achieve a correct understanding. For example, ‘Tom usually takes a left turn on the main road to get to work. Little did Tom know that, this morning, the left section of the main road was under construction.’ “False photo” stories involved interpreting false or outdated content in a photograph. For example, an old photograph depicts the Berlin wall, but no longer accurately describes the current state. “Questions” were structured as true/false statements; ‘true/false’ keywords were displayed on-screen. For a statement like ‘Tom makes a right turn on the main road to avoid the construction’, participants responded by pressing a button to indicate their answer, either true or false.

### MRI data acquisition

All fMRI data were acquired on a 3T Siemens MAGNETOM Prisma MRI scanner with 32-channel parallel imaging at the Dartmouth Brain Imaging Center at Dartmouth College. Structural images were acquired using high-resolution T1 spoiled gradient recall images and were used for anatomical localization and warping to the standard Montreal Neurological Institute (MNI) space only. Functional images were acquired with a multiband EPI sequence (repetition time = 460 ms, echo time=27.2ms, field of view =220mm, multiband acceleration factor = 8, flip angle=44 °, 64x64 matrix, 2.7x2.7x2.7xmm voxels, 56 interleaved ascending slices, phase encoding posterior » anterior). Stimulus presentation and behavioral data acquisition were controlled using Psychtoolbox (MATLAB, MathWorks). Magnetic resonance imaging acquisition parameters are listed in (Table 1).

### 0.1 MRI data curation

MRI protocols in the scanner were renamed to follow the ReproIn naming convention, ensuring robust and automated conversion into BIDS format using HeuDiConv v.0.9.0^24^. For initial sequences not named according to ReproIn, we provided a remapping to facilitate conversion. All subject IDs were anonymized to follow within study sequential order from 1 to 133, left padded with 0 to the length of four digits, e.g. sub-0123. Data was converted to BIDS DataLad dataset using HeuDiConv with dcm2niix v.1.0.20201102^25^, shipped within Singularity container of the ReproNim/containers DataLad dataset^26^ `repronim-reproin-0.9.0.sing`.

### MRI preprocessing

Preprocessing was performed using fMRIPrep 21.0.2^27^ which is based on Nipype 1.6.1^28^;^29^. The following description in this section is generated from the standard fMRIPrep preprocessing boilerplate text to ensure a consistent description across different processing pipelines.

*Preprocessing of B0 inhomogeneity mappings* A *B0*-nonuniformity map (or *fieldmap*) was estimated based on two (or more) echo-planar imaging (EPI) references with topup [30, FSL 6.0.5.1:57b01774].

*Anatomical data preprocessing* A total of 1 T1-weighted (T1w) images were found within the input BIDS dataset. The T1-weighted (T1w) image was corrected for intensity non-uniformity (INU) with `N4BiasFieldCorrection`^31^, distributed with ANTs 2.3.3 [32, RRID:SCR_004757], and used as T1w-reference throughout the workflow. The T1w-reference was then skull-stripped with a *Nipype* implementation of the `antsBrainExtraction.sh` workflow (from ANTs), using OASIS30ANTs as target template. Brain tissue segmentation of cerebrospinal fluid (CSF), white-matter (WM) and gray-matter (GM) was performed on the brain-extracted T1w using `fast` [33, FSL 6.0.5.1:57b01774, RRID:SCR_002823]. Brain surfaces were reconstructed using `recon-all` [34, FreeSurfer 6.0.1,RRID:SCR_001847], and the brain mask estimated previously was refined with a custom variation of the method to reconcile ANTs-derived and FreeSurfer-derived segmentations of the cortical gray-matter of Mindboggle [35, RRID:SCR_002438]. Volume-based spatial normalization to two standard spaces (MNI152NLin2009cAsym, MNI152NLin6Asym) was performed through nonlinear registration with `antsRegistration` (ANTs 2.3.3), using brain-extracted versions of both T1w reference and the T1w template. The following templates were selected for spatial normalization: *ICBM 152 Nonlinear Asymmetrical template version 2009c* [mni152nlin2009casym, RRID:SCR_008796; TemplateFlow ID: MNI152NLin2009cAsym], *FSL’s MNI ICBM 152 non-linear 6th Generation Asymmetric Average Brain Stereotaxic Registration Model*^36^, RRID:SCR_002823; TemplateFlow ID: MNI152NLin6Asym].

*Functional data preprocessing* For each of the 41 BOLD runs found per subject (across all tasks and sessions), the following preprocessing was performed. First, a reference volume and its skull-stripped version were generated by aligning and averaging 1 single-band references (SBRefs). Head-motion parameters with respect to the BOLD reference (transformation matrices, and six corresponding rotation and translation parameters) are estimated before any spatiotemporal filtering using `mcflirt` [37, FSL 6.0.5.1:57b01774,]. The estimated *fieldmap* was then aligned with rigid-registration to the target EPI (echo-planar imaging) reference run. The field coefficients were mapped on to the reference EPI using the transform. The BOLD reference was then co-registered to the T1w reference using `bbregister` (FreeSurfer) which implements boundary-based registration^38^. Co-registration was configured with nine degrees of freedom to account for distortions remaining in the BOLD reference. First, a reference volume and its skull-stripped version were generated using a custom methodology of *fMRIPrep*. Several confounding time-series were calculated based on the *preprocessed BOLD*: framewise displacement (FD), DVARS and three region-wise global signals. FD was computed using two formulations following Power (absolute sum of relative motions^39^) and Jenkinson (relative root mean square displacement between affines^37^). FD and DVARS are calculated for each functional run, both using their implementations in *Nipype* following the definitions by Powers and colleagues^39^. The three global signals are extracted within the CSF, the WM, and the whole-brain masks. Additionally, a set of physiological regressors were extracted to allow for component-based noise correction *CompCor*^40^. Principal components are estimated after high-pass filtering the *preprocessed BOLD* time-series (using a discrete cosine filter with 128s cut-off) for the two *CompCor* variants: temporal (tCompCor) and anatomical (aCompCor). tCompCor components are then calculated from the top 2% variable voxels within the brain mask. For aCompCor, three probabilistic masks (CSF, WM and combined CSF+WM) are generated in anatomical space. The implementation differs from that of Behzadi et al. in that instead of eroding the masks by 2 pixels on BOLD space, the aCompCor masks are subtracted a mask of pixels that likely contain a volume fraction of GM. This mask is obtained by dilating a GM mask extracted from the FreeSurfer’s *aseg* segmentation, and it ensures components are not extracted from voxels containing a minimal fraction of GM. Finally, these masks are resampled into BOLD space and binarized by thresholding at 0.99 (as in the original implementation). Components are also calculated separately within the WM and CSF masks. For each CompCor decomposition, the *k* components with the largest singular values are retained, such that the retained components’ time series are sufficient to explain 50 percent of variance across the nuisance mask (CSF, WM, combined, or temporal). The remaining components are dropped from consideration. The head-motion estimates calculated in the correction step were also placed within the corresponding confounds file. The confound time series derived from head motion estimates and global signals were expanded with the inclusion of temporal derivatives and quadratic terms for each^41^. Frames that exceeded a threshold of 0.9 mm FD or 1.5 standardised DVARS were annotated as motion outliers. The BOLD time-series were resampled into standard space, generating a *preprocessed BOLD run in MNI152NLin2009cAsym space*. First, a reference volume and its skull-stripped version were generated using a custom methodology of *fMRIPrep*. The BOLD time-series were resampled onto the following surfaces (FreeSurfer reconstruction nomenclature): *fsaverage*. *Grayordinates* files^42^ containing 91k samples were also generated using the highest-resolution `fsaverage` as intermediate standardized surface space. All resamplings can be performed with *a single interpolation step* by composing all the pertinent transformations (i.e. head-motion transform matrices, susceptibility distortion correction when available, and co-registrations to anatomical and output spaces). Gridded (volumetric) resamplings were performed using `antsApplyTransforms` (ANTs), configured with Lanczos interpolation to minimize the smoothing effects of other kernels^43^. Non-gridded (surface) resamplings were performed using `mri_vol2surf` (FreeSurfer).

### Pre-session questionnaires

To connect neuroimaging and behavioral parameters to general psychosocial characteristics, we administered a battery of questionnaires prior to participants’ first visit (Table 2).

### Task instructions

During the behavioral instruction phase, we ensured that participant were informed of the same instructions, despite being introduced to different individuals. Therefore, scripted dialogues were provided to experimenters to be verbally read aloud (hosted on: https://github.com/spatialtopology/calibrate/tree/main/dialogue). This was accompanied by visual aids presented on a monitor in front of participants. Afterwards, participants would complete practice tasks, which are located in github repositories (align videos; self referential; https://github.com/spatialtopology/fractional_factorials/blob/master/RUN_practice.m)

#### Scan setup

##### Audio Calibration

To ensure that auditory stimuli delivered during the experimental tasks would be presented at a volume that was perceptible to the participant, prior to each scan session, participants completed a brief audio calibration task. During initial structural scans (i.e., scout, fmap), a video clip was played and the participant provided a rating to indicate whether the volume was set to a satisfactory level. After the audio calibration task was complete, the experimenters verbally confirmed with the participant that the final volume level was adequate.

##### Arm Measurement Procedures for Application of Thermal Device

Two arm sites were identified for the administration of thermal stimuli during the multimodal negative affect `“task-social”` task. To ensure the same sites were used reliably across sessions, we developed an arm measurement protocol. First, the experimenter took an initial measurement along the midline of the participants left arm: from the crease of the elbow to the crease of the wrist. Using this baseline measurement, the experimenter marked two primary sites, located 1/3 and 2/3 of the distance from the elbow to the wrist. The thermode was applied to these sites and the participant completed a pain screening procedure to verify that they could tolerate the noxious stimuli used during the study. If participants reported hypersensitivity or hyposensitivity on either of the primary arm sites, a secondary site was used, 3 cm below the primary site locations. If hypersensitivity or hyposensitivity was also reported at the secondary arm sites, then the participant was removed from the experiment.

### Preprocessing computing environment

The BIDS-formatted data was analyzed at the Johns Hopkins University Joint High Performance Computing Exchange, a high-performance cluster running CentOS Linux 7.9 and the Sun Grid Engine scheduler. On the Exchange, software packages are made accessible via `lmod`^44^. For this dataset, job submissions were managed with `targets`^45^, which interfaces with the scheduler via `batchtools`^46^. These packages were tracked via `renv`^47^. Each job consisted of a call to a Singularity image^48, 49^ which implemented one of the analyses described above (e.g., fMRIPrep, MRIQC). The derivatives of these apps were tracked by `git-annex` and DataLad^50^.

### Technical Validation

For quality control, we used MRIQC BIDS-App^51^ and extracted image quality metrics (IQMs). To demonstrate quality of the data, we focus on temporal signal to noise ratio (tSNR) and framewise displacement (FD). For the tSNR, per run, we calculate the mean image and temporal standard deviation image and divide the two. The mean image was generated from fMRIPrep, the standard deviation map was generated using `fslmaths`. These tSNR maps are aggregated to the participant level (Fig. 4a). The tSNR maps are fairly homogenous across tasks, so we have plotted the median tSNR map from the `task-alignvideo` as a reference (Fig. 4b). In addition, we extract the framewise displacement values per run, average within-participant, and aggregate it to the group level (Fig. 4c). Each task’s median FD value is smaller than .2 mm, and comparable or smaller than that when compared to the FD values of a large scale dataset, UKbiobank^52^.

**Figure 4.**
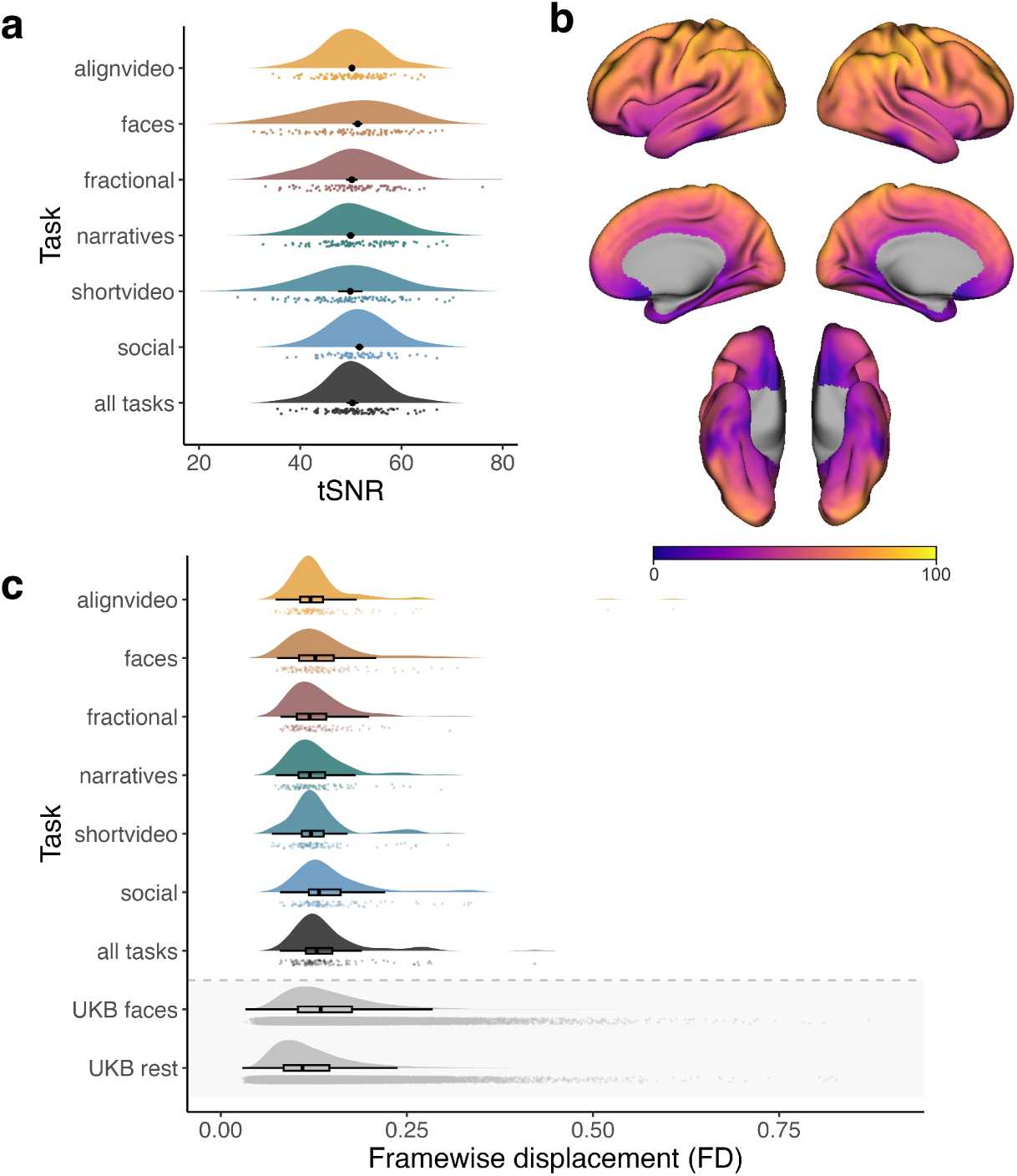
Image quality metrics. (a) Temporal signal to noise ratio (tSNR) was calculated within-subject runs and aggregated across participants. Each density plot represents the distribution of tSNR median values across participants, also reflected in the task-corresponding color markers. The black color markers indicate the group average median tSNR value, calculated across within-mask voxels. The black error bars represent the within-subjects confidence interval for the median tSNR value; Given that short video has one run in the experiment, the error bar reflects the between-subject confidence interval. As indicated in the plot, median tSNR confidence intervals shrink as a function of repeated runs (alignvideos 13 runs; faces 3 runs; fractional 2 runs; narratives 4 runs; shortvideo 1 run; social 18 runs; all tasks 41 runs). **(b)** tSNR of align video task across the cortical surface. Colors indicate median tSNR per voxel. tSNR maps were computed using the preprocessed MNI runwise maps. Groupwise median tSNR maps were converted to fsaverage6 space, using neuromaps, for visualization. **(c)** Head movement is kept to a minimum, reflected in framewise-displacement (FD) values. Each density plot represents the distribution of FD values across participants, also reflected in the task-corresponding color markers. The box plot represents the median value and interquartile range of the group level FD distribution. UKbiobank (UKB) FD values are displayed here for comparison; Across all tasks, the current dataset FD values are lower and less variable compared to that of the “UKB faces” task. “UKB rest” condition has a slightly lower median as it serves as a no task condition compared to the tasks in our current dataset. Overall, head movement was kept to a minimum, as demonstrated by framewise displacement lower than that of the UK biobank, in spite of the longer scan protocol.

### Code availability

Code for stimulus presentation and data acquisition is published in a GitHub repository (https://github.com/spatialtopology). To ensure reproducibility of the experimental procedures, we released the code as version 1, prior to data collection (“task-alignvideo”: https://github.com/spatialtopology/alignvideos/releases/tag/v1.0.0-stable, “task-social”: https://github.com/spatialtopology/social_influence/releases/tag/v1.0.0-stable, “task-faces”: https://github.com/spatialtopology/faces/releases/tag/0.9, “task-narratives”: https://github.com/spatialtopology/narratives/releases/tag/v.1.0.0, “task-shortvideo”: https://github.com/spatialtopology/shortvideos/releases/tag/v1.0.0-stable, “task-fractional”: https://github.com/spatialtopology/fractional_factorials/releases/tag/v.1.0.0). Code for data wrangling and analyzing the data is in the GitHub repository (https://github.com/spatialtopology/spacetop-prep). Neuroimaging data and behavioral data are BIDS-formatted in DataLad^53^. Code for data wrangling and analyzing the data is in the github repository (https://github.com/spatialtopology/spacetop-prep). Neuroimaging data and behavioral data are BIDS-formatted in DataLad.

## Acknowledgements

We would like to express our gratitude to the following individuals and institutions for their invaluable contributions to this study: Elizabeth C. Tremmel for the insightful comments, Terry J. Sackett and Courtney Rogers for the technical support in data collection; Chandana Kodiweera for the valuable input on MR quality control. This data collection was funded by NIBIB:R01EB026549-03.

## Author contributions statement

T.W., H.J., P.A.K., X.H. M.L. conceived the experiment(s). H.J., P.A.K., X.H., M.S., M.O.H. experimental setup H.J., M.A., B.J.H., E.I.M. B.P. conducted the experiment H.J., Y.O.H., Z.M data organization P.S., H.J., Y.O.H., B.P. data preprocessing H.J., Z.M., O.G.C. data quality control and QC analysis All authors reviewed the manuscript.

## Competing interests

(mandatory statement)

The corresponding author is responsible for providing a [competing interests statement](https://www.nature.com/sdata/policies/editorial-and-publishing-policies#competing) on behalf of all authors of the paper. This statement must be included in the submitted article file.

## Notes

### Competing Interest Statement

The authors have declared no competing interest.

